# Butterflies Are Shrinking: Evidence from the Past Century

**DOI:** 10.1101/2025.02.05.636603

**Authors:** Konstantina Zografou, Eva Knop, Brent J Sewall, George C Adamidis, Oliver Schweiger, Theodoros Semertzidis, Panagiotis Stalidis, Consuelo M De Moraes, Mark C Mescher, Melissa RL Whitaker, Michael Greeff, Konstantinos M Anagnostellis, Marina Brokaki, Eleftheria Kaltsouni, Maria Dimaki, Vassiliki Kati

**Affiliations:** Department of Biological Applications and Technology, University of Ioannina, 45100 Ioannina, Greece; Agroscope, Reckenholzstrasse 191, 8046 Zurich; Department of Biology, Temple University, Philadelphia, Pennsylvania, USA; Laboratory of Plant Physiology, Department of Biology, University of Patras, 26500 Patras, Greece; Helmholtz Centre for Environmental Research - UFZ, Department Community Ecology, 06120 Halle, Germany; The Visual Computing Lab, Information Technologies Institute/Centre for Research and Technology Hellas, 6^th^ km Charilaou-Thermi Road, Thessaloniki, GR-57001, Greece; Department of Environmental Systems Science, Swiss Federal Institute of Technology (ETH Zürich), 8092 Zürich, Switzerland; Biological Sciences, East Tennessee State University, Johnson City, TN, United States; Department of Terrestrial Zoology, Goulandris Natural History Museum, 100, Othonos ST. Kifisia, Greece

**Keywords:** butterflies, body size, shrinkage, forewing length, global warming, museum specimens, computer vision analysis, terrestrial biomes, digitized collections

## Abstract

Climate change has had strong impacts on biodiversity, including well-documented shifts in the distributions and phenology of species. Reductions in body size represent a third pervasive biological response; it has garnered significantly less attention despite great ecological relevance. Theoretical frameworks-the temperature-size rule, Bergmann’s rule, and James’s rule-predict that warmer temperatures are associated with smaller body sizes in ectotherms. In contrast, empirical evidence concerning this pattern is mixed across both taxa and environments. In the present study, we applied computer vision techniques to historical data from two large butterfly museum collections, totaling 593 species across 10 terrestrial biomes over more than a century, in order to investigate long-term trends in butterfly body size. We measured forewing length through both manual image analyses and automated computer vision algorithms proxying body size and analyzed trends by using generalized additive models in order to consider a temporal, biome-specific pattern assessment. We have tested two hypotheses: that butterfly body size (1) declines over time, in conjunction with increasing ambient temperatures, in agreement with the temperature-size rule, and (2) its variation is more marked in warm, dry biomes. Results indicate a significant overall reduction in the body size of butterflies during the last century and that this reduction is indeed more pronounced in those biomes facing higher rises in temperature. These findings constitute large-scale evidence in support of the temperature-size rule and indicate a potential ecological impact of climate change on butterfly populations.

## Introduction

Climate change has had dramatic effects on biodiversity, with well-documented shifts in species distributions and phenology in response to rising temperatures (Parmesan and Yohe 2003, Thomas et al. 2004, Parmesan 2006). A third widespread biological reaction to increased temperatures, namely shifts in body size, has received relatively little attention, despite its potential ecological significance (Sheridan and Bickford 2011). Body size is a key biological characteristic that controls individual fitness, species interactions, and ecosystem function (Hildrew et al. 2007, Verberk et al. 2021).

Theoretical frameworks explaining temperature-related variation in body size include temperature-size rule (Atkinson 1994), Bergmann’s rule (Bergmann 1847), and James’s rule (James 1970). Temperature-size rule supposes that ectothermic animals will mature at smaller sizes when reared in warmer environments, possibly facilitated through phenotypic plasticity during ontogeny. In contrast, both Bergmann’s and James’s rules refer to larger-scale trends, suggesting that species or populations have smaller body dimensions in warmer geographical locations or over long evolutionary timespans.

Proposed mechanisms for these trends involve both biophysiological consequences of temperature acting on development and growth rates (Ghosh et al. 2013) and physiological constraints with regard to metabolite requirements and availability (Verberk et al. 2021). For instance, high temperatures can heighten metabolite demand, and in consequence, restrict growth duration before maturation is attained. In addition, factors like oxygen limitation and food resource constraints may further mediate body size responses (Queiros et al. 2024). In the case of butterflies, which provide important ecosystem services through their pollination activities, scientists investigated how temperature-mediated shrinkage could affect butterflies’ pollination role. They found that the warmest reared butterflies carried less pollen and from fewer plant species (Büyükyilmaz and Tseng 2022).

Empirical research has produced mixed evidence for these theoretical predictions. Despite documented reductions in arctic butterflies (Daly et al. 2024), several aquatic species (Daufresne et al. 2009), tropical moths (Wu et al. 2019), Hymenoptera (Polidori et al. 2020, Barrett and Johnson 2023), Diptera (Baranov et al. 2021), and damselflies (Hassall 2013), all in agreement with temperature-size predictions, several studies have documented contrasting trends or no significant trends. Examples of increased body size include temperate butterflies (Davies 2019, Wilson et al. 2019, Wilson et al. 2023), the common lizard species *Lacerta vivipara* (Massot et al. 2006), and mammals such as *Martes americana* in Alaska (Yom-Tov et al. 2008) and *Lutra lutra* in Norway (Yom-Tov et al. 2006). These discrepancies mean that additional ecological and physiological factors, such as life-cycle complexity, sexual dimorphism, food quality, and competition between species, can modulate body size responses to warming temperatures (Chown and Gaston 2010, Wetzel et al. 2021).

Despite the growing body of literature, it remains unclear whether body size reductions can be considered a universal response to climate warming. To fill this information gap, studies with high taxonomic, geographical, and temporal scopes must be performed to disentangle the impact of climate change from additional factors in the environment. Such in-depth studies have long been challenging in view of data availability constraints. With an increased availability of digitized collections in museums, in addition to improvements in image analysis technology, such in-depth studies can, at present, assess long-term biological trends with unprecedented opportunity (Høye et al. 2021, Groom et al. 2023).

In the current study, we have leveraged collections in museums and utilized computer vision techniques to analyze trends in butterfly body size in relation to warming trends over time. The MEIOSIS project holds two large butterfly collections, representing 593 species in 10 terrestrial biomes over a long period exceeding one hundred years. In our analysis, we have considered the following hypotheses: (1) a reduction in butterfly body size over time in proportion to an elevation in ambient temperatures, in agreement with the temperature-size hypothesis for ectotherms (Atkinson 1994); and (2) increased variation in body size in biomes with warmer and drier environments, and therefore directly impacted by temperature increases worldwide (Li et al. 2018). By analyzing trends in butterfly body size in a variety of biomes, our intention is to gain a thorough understanding of the ecological consequences of climate change in butterfly populations.

## Methods

### Entomological collections (metadata)

We assessed preserved specimens from two entomological collections in Europe. The Entomological Collection of the Swiss Federal Institute of Technology in Zürich (ETHZ hereafter) comprises a core Palaearctic Macrolepidoptera collection, and is the fourth largest entomological collection in Switzerland. The ETHZ metadata collection provides information for 48,766 butterfly specimens belonging to five families (Pieridae, Lycaenidae, Papilionidae, Nymphalidae, Hesperiidae) and 639 species, collected between 1028 to 2019. All digitized records of palearctic microlepidoptera specimens are accessible through an online database NAHIMA (https://www.nahima.ethz.ch/pool/entomologische_sammlung).

We also use specimen record from the Goulandris Natural History Museum (GNHM hereafter) zoological collection. Because these specimens were not already digitized, we initiated a digitization process similar to that of ETHZ, enabling us to merge both collections into a global database. We established two imaging stations in GNHM consisting of a digital camera with ED 40mm f/2.8 Macro focal lens mounted to a Kaiser camera stand and two RB 218N HF LED table lamps (5400 Kelvin). A team of three researchers and two students photographed 6,639 physical specimens belonging to 274 species collected between 1917 to 2023. The team was trained by a professional photographer to standardize imaging processes and followed a set of standardized protocols to minimize the risk of potential mistakes. When needed, the team repaired closed or loose wings before imaging and excluded specimens with incomplete data or specimens that could not be repaired, from the final database. Excluding specimens with incomplete information (e.g., collection location or date), a total of 44,316 specimens from ETHZ and 4,582 specimens from GNHM metadata were retained.

### Manual measurements (ImageJ)

We manually measured the right forewing length from 22,807 images. For the ETHZ collection, a subset of 17,382 specimens (used to validate an automated measurement algorithm, see below) was analyzed, while for the GNHM collection, 5,425 specimens were measured after excluding those with damaged wings. After merging these manual measurements with the metadata and removing duplicates, 17,024 specimens from ETHZ and 3,999 specimens from GNHM were retained. For pinned butterfly specimens, forewing length is the best-preserved trait that is strongly correlated with overall body size and has been previously used as a proxy for overall body size (Brehm et al. 2019, Minter et al. 2024). We defined the forewing length as starting from the insertion point connecting the thorax up to the wing tip (Wu et al. 2019), excluding the scales at the very edge, as they were often absent. First, we calibrated the scale bar embedded in each specimen image to measure a known distance as for example 10 mm and then we measured forewing length which was converted to millimeters using the scale bar and the “draw line” function in Image J (https://imagej.nih.gov/ij/index.html).

### Automated measurements (algorithm)

For automated morphological measurement of the samples, a custom computer vision tool was implemented. The need for such a customization was because the existing state-of-the-art tool, named Mothra (Wilson et al. 2023), supported only a very specific sample image configuration that is not covering all the different configurations of the MEIOSIS project datasets.

The automated, algorithmic measuring follows a two-step approach to calculate forewing length. In the first step, a pretrained model of the state-of-the-art “segment anything” (Kirillov et al. 2023) semantic image segmentation algorithm was used, to separate the sample from the background, select the right forewing, and to extract other image components such as the ruler. A key functionality of the segment anything model, is the ability to break an object into its semantic parts and deliver both the object as a whole, and its parts, as hierarchical segmentation levels. Having the image hierarchically segmented, a vision classifier was trained as ruler detector and right forewing detector, based on a ResNet-18 (He et al. 2016) model architecture. For the ruler, the scale calculation was provided by traditional frequency-domain analysis.

After isolating the contour (foreground-only) of the right forewing image and by following example annotations from the researchers, the second step of the approach was to implement a rule-based computer vision algorithm that follows the contour shape to measure the leftmost and rightmost active pixels of the right forewing. As a corrective step, the position of antennae was also computed to identify the right-wing to body, shoulder position.

The computer vision tool was implemented in Python 3, it requires a GPU-enabled server to operate on a convenient speed of approximately 1.5 seconds per image (depending on image size), and is fully customizable to support other morphological measurement needs, also for other species beyond butterflies.

The algorithm processed the first collection (ETHZ), analyzing 44,316 specimens. However, it failed to converge for 1,116 cases. We then manually reviewed the remaining 43,200 images, excluding specimens that (i) had invalid measurements (e.g., forewing length extending beyond the wing tip) and (ii) were pinned ventrally instead of dorsally. After applying the first criterion, 28,860 specimens remained, while 27,058 met both conditions and were retained and merged with ETHZs’ metadata. After removing duplicates, a total of 23,265 unique specimens remained.

### Merging forewing length measurements

To increase the robustness of our sample size, we combined manual and automated measurements. First, we tested the accuracy of the algorithm: we compared the common specimens for both approaches using Spearman rank correlation. Then we considered a linear relationship (forewing_algorithm_ = intercept + slope*forewing_ImageJ_) to translate all forewing lengths measured manually with ImageJ to that of the algorithm output. After merging manual and automated forewing measurements from both collections, a total of 31,920 unique specimens corresponding to 593 species across 88 countries, collected between 1028 to 2019 were retained for further analysis. We further excluded periods between 1028 – 1840 and 2010 – 2019 where specimens were scarce (6 and 15 specimens accordingly) and the period 1840-1899 (1500 specimens) as specimens were present only in the half of the studied biomes (5/10).

### Biomes

To investigate our second hypothesis, we utilized the global terrestrial biome map developed by Olson et al. (2001) by spatially joining biome attributes to each specimen. This map represents 14 biomes that encompass all terrestrial ecoregions and is largely regarded as a useful tool for conservation management. On the global scale, biome distribution can be determined by the climate, a relationship well-demonstrated by the Köppen–Geiger climate classification system (Rohli et al. 2015). This system’s effectiveness is evident, as many biomes such as deserts, tundra, and tropical rainforest, occur almost exclusively in specific types of climate because key factors-temperature and precipitation patterns-control it. Our dataset consisted of specimens from 13 biomes; three biomes had less than 20 specimens and thus were excluded for further analyses (**Figure 1, Table S1**).

**Figure 1.**
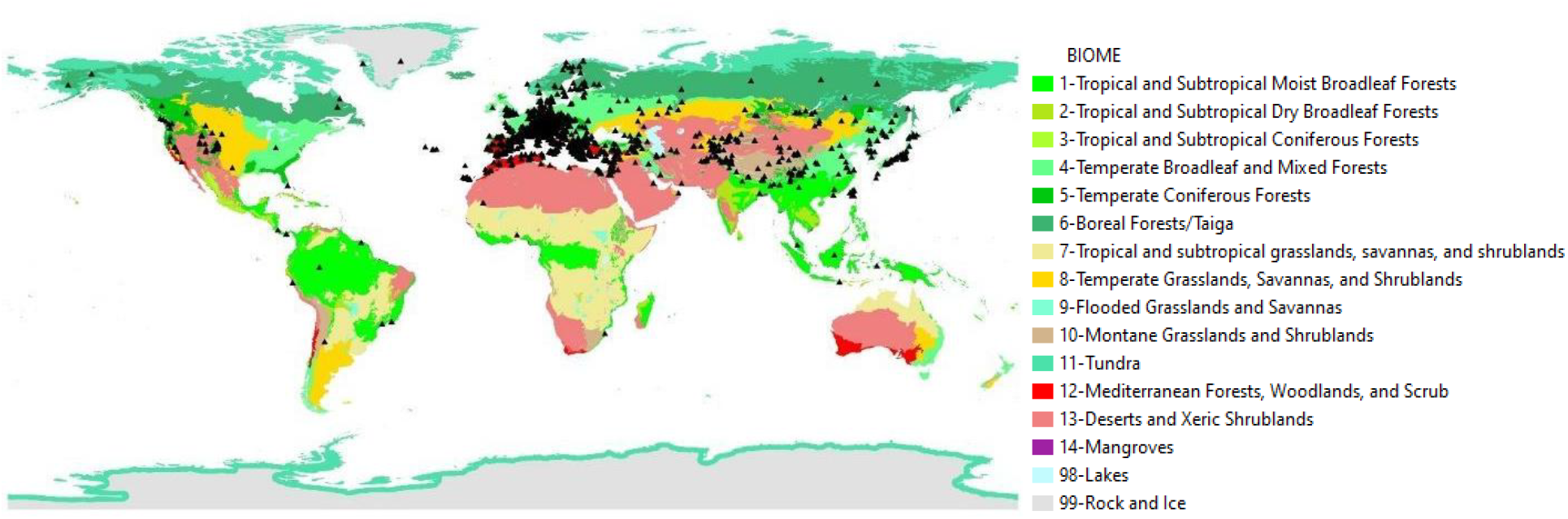
Fourteen biomes according to Olson et al. (2001). Our study system is represented by black triangles (31,920 specimens, 593 species, 10 biomes, 88 countries).

### Data analysis

To investigate our first biological hypothesis—whether there are any temporal trends in the forewing length of our butterfly pool—we employed a Generalized Additive Model (GAM). The GAM framework was selected because data exploration confirmed our expectation that the relationship between forewing length (the response variable) and year is non-linear. We modeled species forewing length using a Gaussian error distribution with an identity link function, appropriate for continuous data. In terms of fixed effects, we included biome as a parametric term to account for differences among biome categories. The primary smooth term was a thin plate regression spline (TPRS) applied to the year variable, capturing non-linear trends in forewing length over time. Regarding random effects, we were not specifically interested in species-level variation per se, but we accounted for it to control for the fact that forewing lengths from the same species are likely more similar than those from different species.

We specified two random effects: a random intercept for species to model species-specific baseline differences in forewing length and a nested random effect of species within years to capture any additional variation driven by differences in species composition across annual collections. The full model structure can be expressed as:

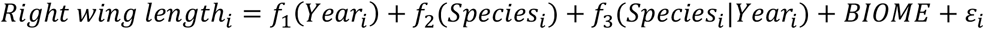

where *f*_1_(*Year*_*i*_) is the smooth term for the continuous predictor, *f*_2_(*Species*_*i*_) is the random effect for species, *f*_3_(*Species*_*i*_ |*Year*_*i*_) is the nested random effect of species within years, *BIOME* is the parametric term for the different categories of biomes and *ε*_*i*_ is a Gaussian error term. We used model with one global smoother as described in Pedersen et al. (2019) with the mgcv package (Wood 2023) and REML smoothness selection (Wood 2011, Wood 2017) in R version 4.3.2 (R Core Team 2024) and diagnostic plots with *appraise ()* function from gratia package (Simpson 2024).

For our second hypothesis, we considered again Generalized Additive Model (GAM) using a Gaussian error distribution with an identity link function, without the global smoother (Pedersen et al. 2019). In this model, the underlying assumption is that, despite any similarities in the shape of the functions, group-level smooth terms do not share or diverge from a common (global) form. As we expected that temperature influence in biomes will differ in time, we allowed each group-specific smoother a different smoothing parameter, suggesting thus a different level of wiggliness. We fitted a model with a separate smoother, a random effect of species to model species-specific intercepts, a nested random effect of species within years and the parametric term of biomes for the post-hoc comparison of the contrasts using the following formula:

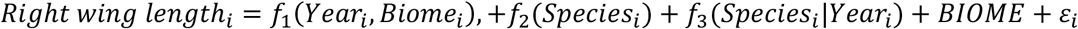

where *f*_1_(*Year*_*i*_, *Biome*_*i*_), was the smoothed interaction between the continuous predictor and the grouping factor allowing the effect of year to vary by biome, *f*_2_(*Species*_*i*_) was the random effect for species, *f*_3_(*Species*_*i*_ |*Year*_*i*_) was the nested random effect of species within years, *BIOME* the parametric term for the different categories of biomes and *ε*_*i*_ was a Gaussian error term.

We used mgcv package to run our gam models *wald_gam ()* function from itsadug package (van Rij J et al. 2022) for post hoc comparison of the intercept differences. Plots were made with *marginaleffects* package (Vincent Arel-Bundock et al.) and gratia package (Simpson 2024), fit of the model with *gam*.*check ()* function from mgcv package (Augustin et al. 2012) and diagnostic plots with *appraise ()* function from gratia package (Simpson 2024).

### Species Body Sexual dimorphism

To account for body sexual dimorphism effect, we rerun each model using only species for which the wing length was known to be the same between females and males. We used the European and Maghreb butterfly trait database (Middleton-Welling et al. 2020) and the morphological trait “Forewing length (FoL)” that corresponds to male and female average length with data obtained by various sources. We found no change from full models, and we reported the output in the supporting information file (Body size sexual dimorphism section).

## Results

### ImageJ versus automated measurements

ImageJ and algorithm’s measurements are almost the same (**Figure 2**). Spearman rank correlation between the right forewing length measured manually with ImageJ and right forewing length measured automatically with the algorithm is 0.96 (*P* < 0.001). We were able to translate all measurements with ImageJ to an algorithm output (with intercept = 0.12 and slope 0.99). Diagnostic plots showed no violation in modelling assumptions.

**Figure 2.**
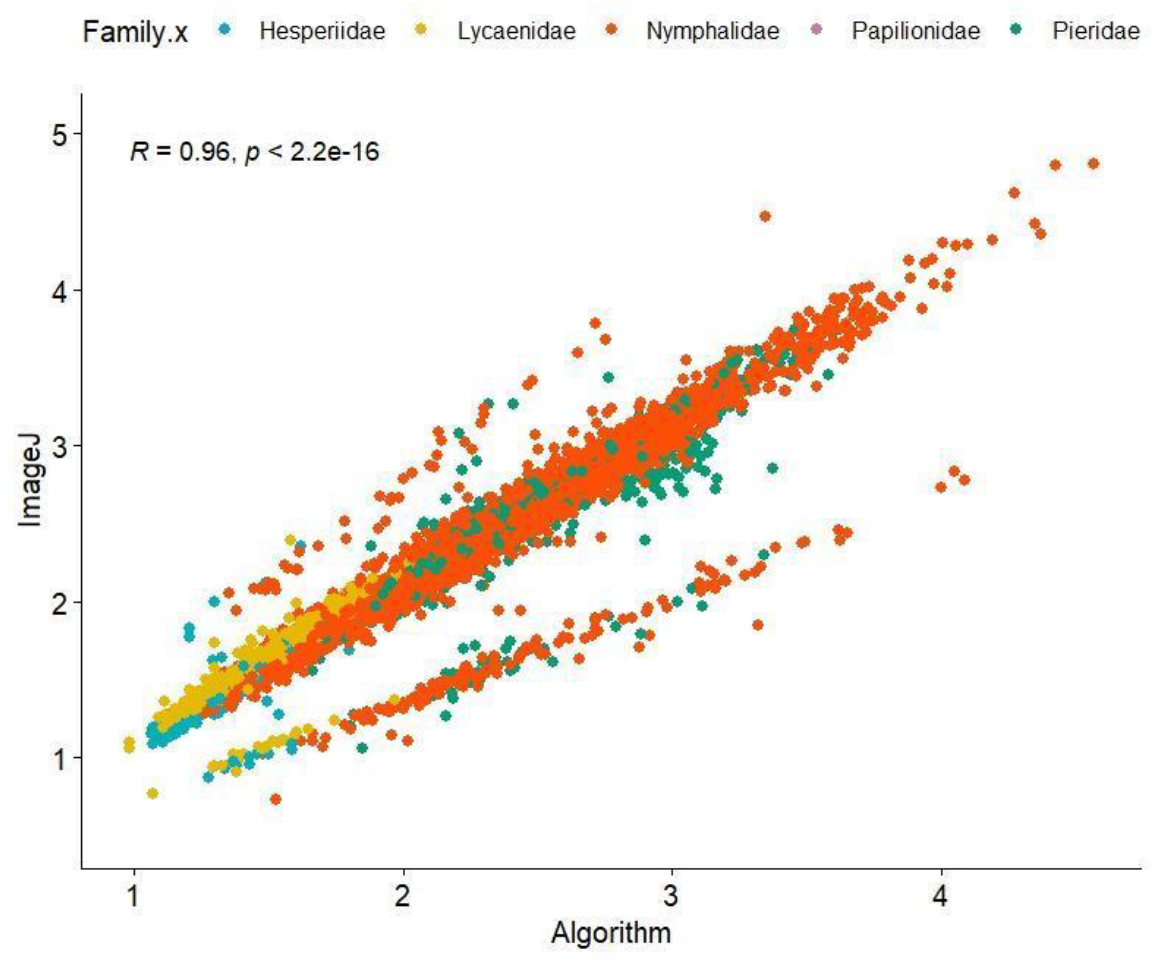
The Spearman rank correlation between the right forewing length measured manually using ImageJ and automatically using the algorithm is 0.96 (*P* < 0.001), demonstrating a strong agreement between the two measurement methods. Different colors correspond to the five families as shown in the upper part of the figure. Both manual and automated measurements are considered in centimeters.

### A single (global) smoother

The estimated global smoother of *Year*_*i*_ (**Figure 3)** was strictly negative, i.e., below zero, after the year 1970, with the decreasing trend starting two decades earlier, the year 1950. A positive effect for the right forewing length between 1920-1950 followed the first decreasing period between 1900 and 1920. However, the confidence intervals between 1900 and 1935 included intercept zero for most of the range of smoother of *Year*_*i*_ indicating that this period had no valid effect. The global smoother with 3.55 estimated degrees of freedom, confirms our expectation for a wiggly trend. Model was fully converged after 6 iterations and diagnostic plots show no violation of model assumptions (**Table S2, Figure S1**).

**Figure 3.**
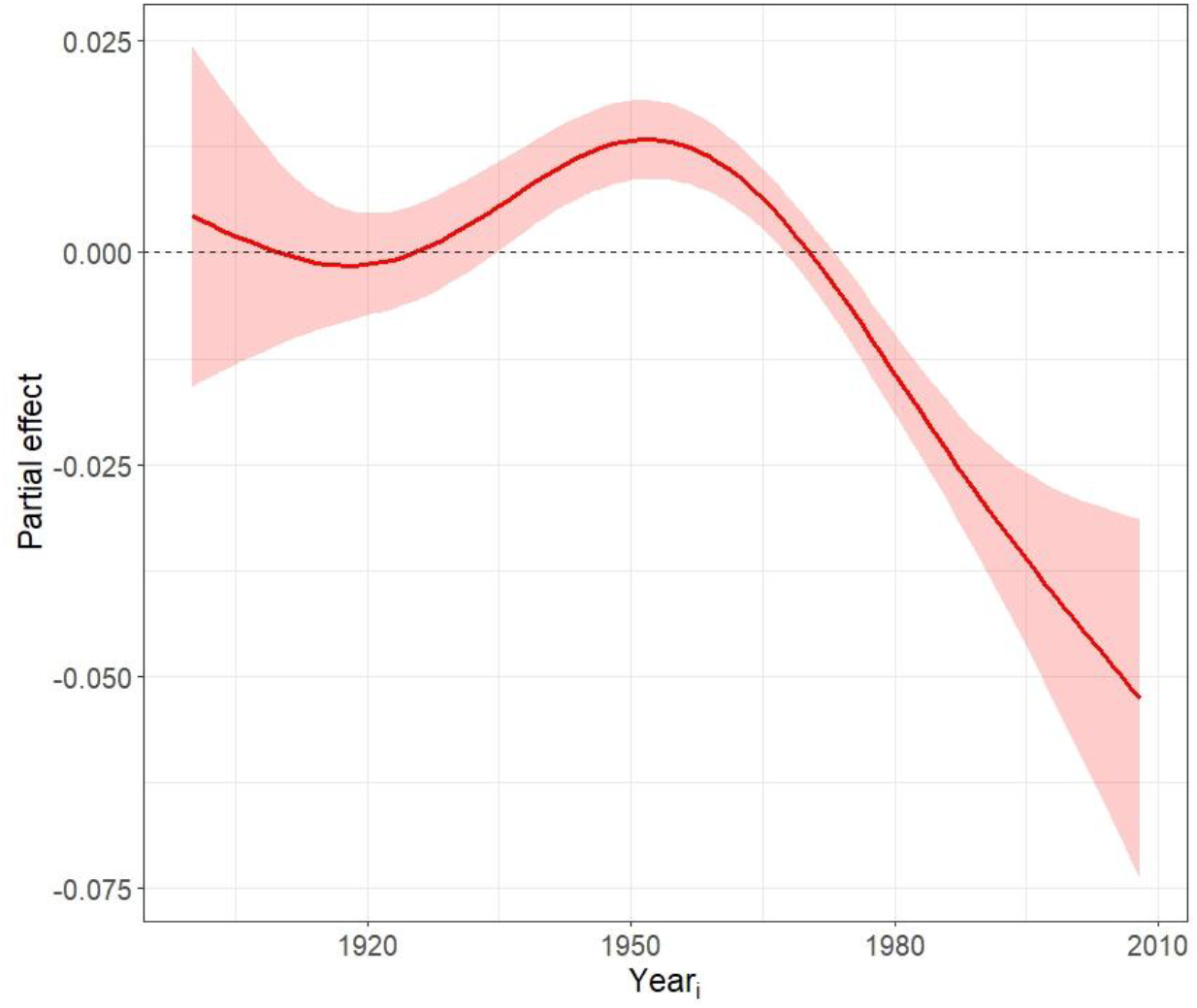
Partial effect plot for the right butterfly forewing length and year smooth. Given that all other variables in the model were set to zero, the plot displays the individual component effect of year smooth function on the link scale (link = identity).

To better understand the shifts of the slope of the fitted function over time we calculated the slope (i.e., the first derivative) of our global smoother using *plot_slopes ()* function (**Figure 4**), which showed that the slope (or trend) was negative or decreasing for most of its range (below zero) and took positive values only for a small fraction of the studied period (1920 – 1955).

**Figure 4.**
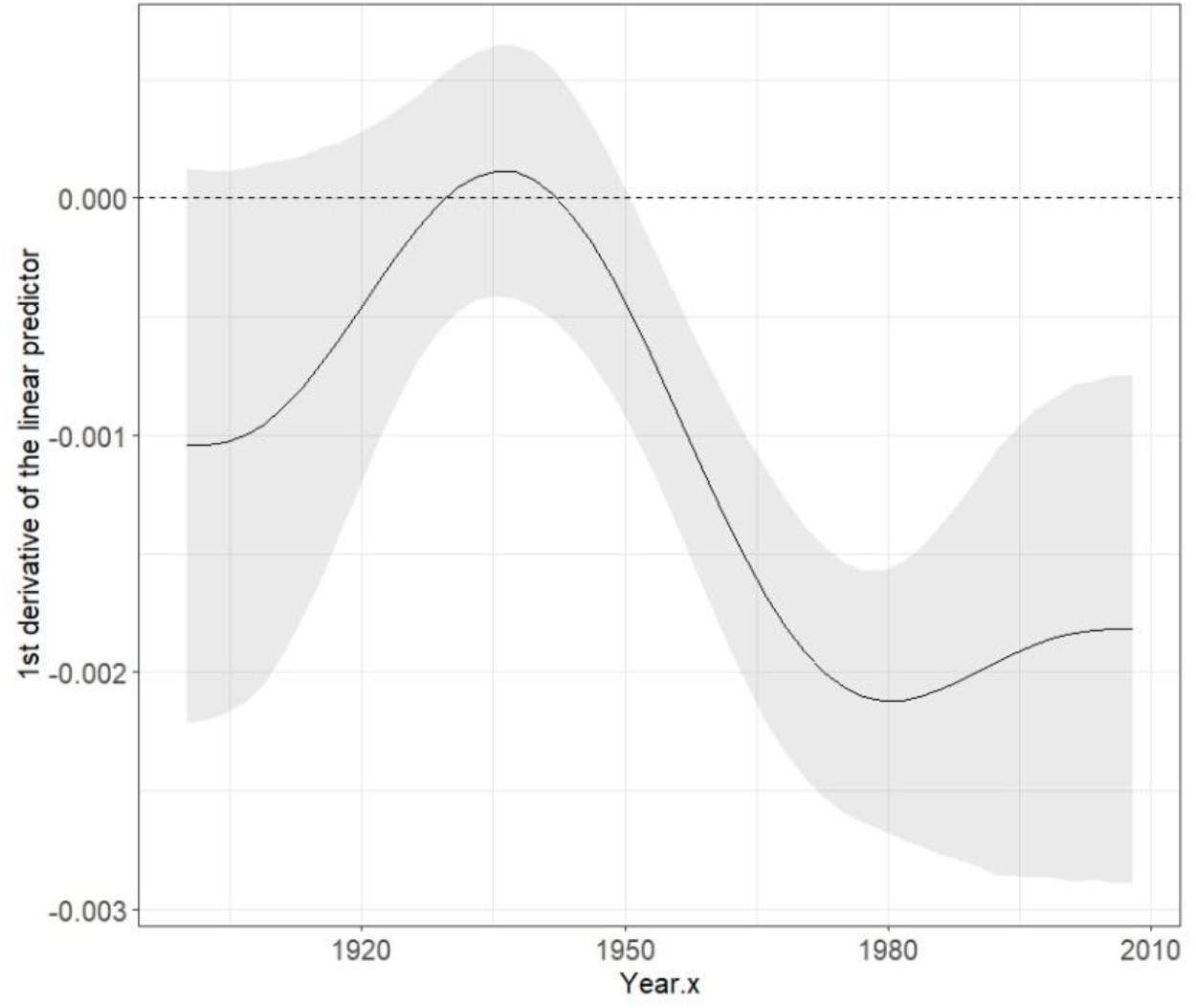
Changes of the fitted function over the years for the first model (slope corresponds to the first derivative).

### Group-specific smoothers for biomes

With the second model, we show that the estimated group-specific smoothers for five biomes (1, 4, 11, 12, 13) significantly differed (**Figure 5, Table S3**). The well fit of the model verified after 10 iterations (full convergence) and diagnostic plots showed no violation from model assumptions (**Figure S2**). Post-hoc comparisons of the contrasts showed significant differences for the five biomes (*P* < 0.001, **Table S4**).

**Figure 5.**
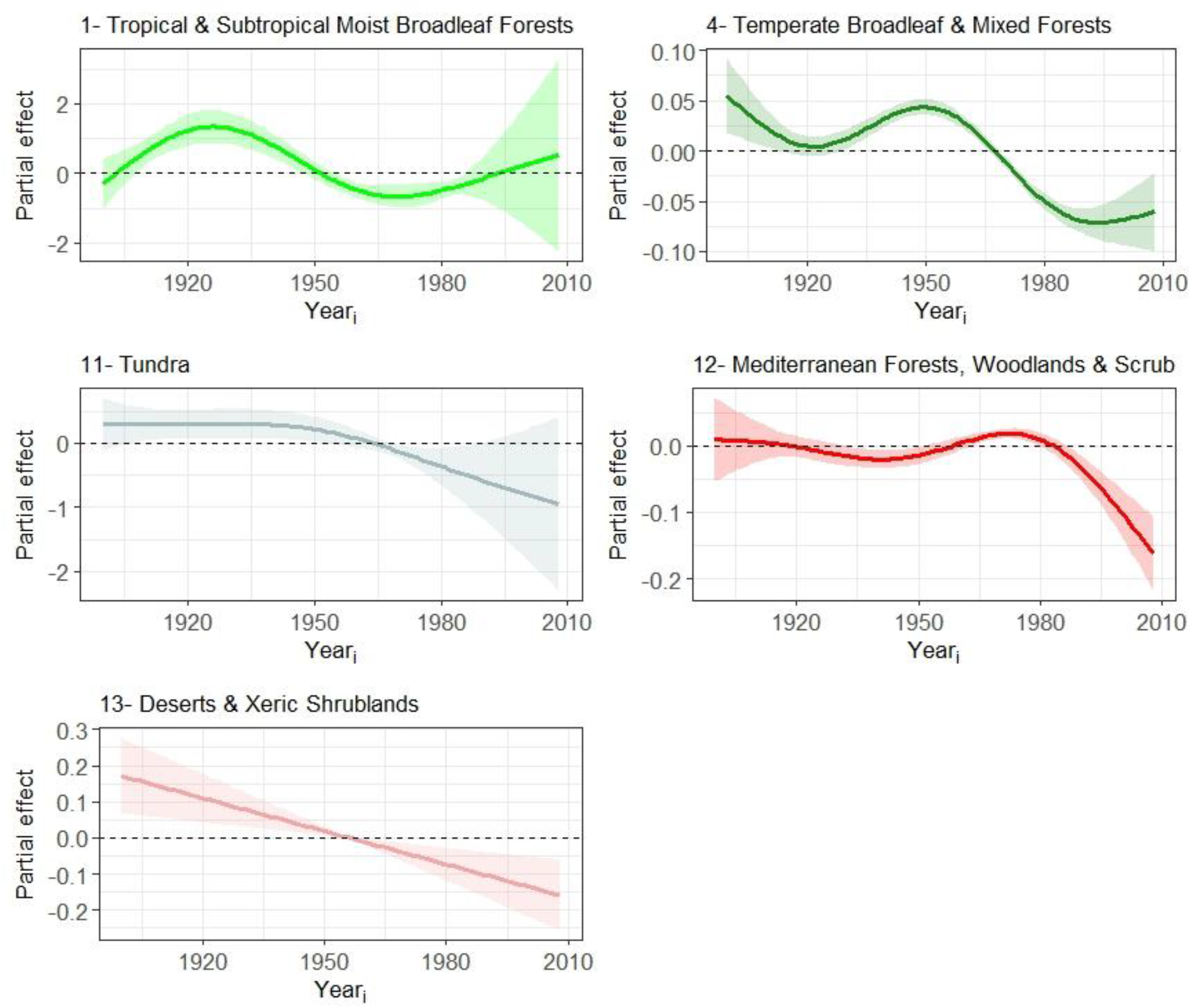
Partial effect plots for the right forewing length and year smooth by biome on the link scale (link = identity). Numbers (1, 4, 11, 12, 13) correspond to different biomes and colors follow the coloring of Figure 1.

Even after aggregating temperate biomes (biomes 4; 5; 8) and tropical biomes (biomes 1;3;7) results remained the same (SI: Aggregating biomes section).

Tropical and Subtropical Moist Broadleaf Forests (biome 1) showed a decreasing trend between 1926 and 1970 (**Figure 5**). On the contrary, for the periods 1900-1926 and 1970-2010 there was a positive trend, which seemed sharper for the first period. Confidence intervals, however, include zero line after 1985 for the entire range of year until the end of the studied period, suggesting that this period had no valid effect. Temperate Broadleaf and Mixed Forests (biome 4) and shows a sharp decreasing trend at the beginning of the studied period (1900-1920) and then again after 1950. In addition, a period of positive trend is spotted for the period between 1925 and 1950 (**Figure 5**). Wide confidence intervals for the final years, after 1990, suggest that the positive trend might not be valid for this period. Tundra (biome 11) first shows a rather stable trend (**Figure 5**) which starts decreasing only after 1950 and until the final year. However, the confidence intervals after 1990 include zero for the rest of the range of smooth of year suggesting perhaps a non-valid trend over this period. Mediterranean Forests, Woodlands, and Scrub (biome 12) showed a steep decrease after 1970 (**Figure 5**). Before that, another period of interest starts near 1930 where for almost two decades we see a more stable trend with a smooth reduction in its middle (1940). From that point and until 1970, the smooth year shows a positive trend. Deserts and Xeric Shrublands (biome 13) showed a steep linear decreasing trend over the years (estimated degree of freedom = 1) (**Figure 5**).

## Discussion

To date, there is no universal evidence supporting the idea of butterfly shrinkage due to global warming. Our study largely confirms the general applicability of temperature-size rule for this taxon. Our results provide evidence that butterflies have reduced their size under warming over the last century (1900-2010). Notably, the sharp decrease we demonstrate after 1950, coincides with the first proof provided for CO_2_ concentrations rise (Keeling 1968, Brewer 2009) and the mid-20^th^ century spread of the Industrial Revolution. Climate change has resulted in elevated global mean surface temperatures by 0.5°C to 1.3 °C for the period 1951-2010 (IPCC 2021), with human climate influences being responsible for 50 to 150 percent of the observed warming between 1950 and 2005 (Wigley and Santer 2013).

Further increase up to 4°C by 2100, unless action is taken to reduce the rate of warming (IPCC 2021), might push butterflies beyond their physiological limits, especially the more sensitive to changes, the ones living in harsh environments and the ones closer to their upper thermal limits (Klockmann et al. 2017).

After the temperature-size rule proposed by Atkinson (1994), a growing number of studies have provided evidence for the shrinking body size of taxa (both ectotherms (Ghosh et al. 2013, Brehm et al. 2019, Baranov et al. 2021, Minter et al. 2024) and endotherms (Masoero et al. 2024, Pirotta et al. 2024)) and for a variety of environments (terrestrial (Polidori et al. 2020) or aquatic (Queiros et al. 2024, Taboada et al. 2024)) from observations at different levels of biological organization such as communities (Wu et al. 2019), individual species (Bristow et al. 2023, Daly et al. 2024, Minter et al.

2024), and/or families (Brehm et al. 2019). Evidence for the opposite trend, i.e., increasing body size is also mounting (Wonglersak et al. 2020, Na et al. 2021, Wilson et al. 2023). Despite the unquestionable value of these studies, most of them limit their ability to assess broad-scale patterns due to geographic and/or time constraints, as well as their species-specific focus. Our study addresses these limitations by analyzing butterfly data from around the globe over a period of more than 100 years.

### Biomes

We examined ten terrestrial biomes and we captured a general decreasing trend for body size over the years. We limit our discussion to the five biomes, with a significant trend (**Figure 5**). We hypothesized that responses across biomes will differ (Gu et al. 2023) as their susceptibility to climate warming differs, and we anticipated responses to be more pronounced in warmer and drier biomes as these are highly exposed to climate warming (Li et al. 2018). Along these lines, the most systematic and continuous decrease in butterflies’ body size is identified in Deserts and Xeric Shrublands, with no evidence of a turnaround for its negative trend whatsoever.

While studies for butterfly body size shrinking is lacking for Deserts and Xeric Shrublands, similar pressures in other arid-adapted insects, such as bees, show evidence of body size shrinkage between 1974 and 2022 (Barrett and Johnson 2023). Deserts and Xeric Shrublands biome is recognized as the most vulnerable terrestrial biome to climate change (Li et al. 2018). As climate change intensifies, we suspect that physiological remodeling of butterflies in such harsh environments is being pushed further (Seebacher et al. 2015). The idea that butterfly physiological plasticity may surpasses a critical threshold beyond which it might be unable to confer resilience to environmental stress, like many plants do (González-Tokman and Wesley 2024) is an area of greatest need for future research.

Similarly to Deserts and Xeric Shrublands, a sharp decline with no reversal of the negative trend was found for Mediterranean Forests, Woodlands, and Scrub after 1970. Mediterranean ecosystems are considered biodiversity hotspots due to their favorable climate conditions and their topographical and vegetation structure complexity (Médail and Quézel 1999) but stressing drivers such as rising temperatures, extended drought, reduced rainfall and increasing number of wildfires make them highly vulnerable and exposed to climate change (Caretta et al. 2022). Forecasts for Mediterranean Basin indicate an even drier and hotter climate over the coming decades making this area a climate hotspot (IPCC 2021). According to global biodiversity forecasts for the year 2100, Mediterranean biomes will undergo the most significant proportionate shift in biodiversity (Sala et al. 2000). Over the past three decades, widespread population declines have been observed in various Mediterranean butterfly species, with these declines strongly linked to the intensifying impacts of global change (Wilson et al. 2007, Zografou et al. 2014, Carnicer et al. 2019). Populations living in areas with no elements for thermal buffering (e.g. semi-open forests) may experience increased negative effects from extreme temperatures leading to distinct plastic responses. For instance, the plastic response of body size of a Mediterranean butterfly (*Pieris napi*) shows a strong reduction in thermally exposed populations where vegetation thermal buffering is also limited due to reduced plant transpiration, and low leaf quality caused by drought and seasonal advance of plant phenology (Carnicer et al. 2019).

Moving on to the Earth’s northernmost continental landmass, in the arctic Tundra biome we see a rather stable trend at the beginning of the last century (1900) that remains until 1940. On the condition that arctic ecosystems are mainly influenced by a single factor, i.e., climate change (Sala et al. 2000), we assign the decreasing trend imposed after 1940 to the elevated region’s average temperature (2-3°C) over the past 50 years (Meredith et al. 2019, Overland et al. 2019). Along these lines, an analysis of three Holarctic butterflies using museum specimens showed a decrease in wingspans between 1971 and 1995, ranging from 0.7 to 5 mm for every 1°C increase of temperature (Daly et al. 2024). Similarly, body size reduction was found for two High-Arctic butterfly species (*Boloria chariclea* and *Colias hecla*) between 1996 and 2013 (Bowden et al. 2015).

Reduced body size of the studied moths along a tropical elevation gradient (Brehm et al. 2019), or body size reduction in tropical moth assemblages (Wu et al. 2019) over time are in accordance with our long negative trend for the Tropical and Subtropical Moist Broadleaf Forests between 1930 and 1970. The thermal adaptation hypothesis suggests that insect populations in warm, aseasonal tropical environments exhibit higher and narrower critical thermal limits (Kaspari et al. 2015). As the yearly surface temperature for the period from 1900 to 1930 indicated to be cooler than average years (1880-2023 (NOAA 2024) we suggest that lower temperatures might justify the positive trend we found between 1900 and 1930. The second observed increase in body size (1970–1985) is likely mediated by the dominance of large-bodied butterflies from the genera *Trogonoptera* and *Ornithoptera*, which constituted 43% of the sample during this period. However, the small sample size (21 individuals) for this time frame imposes caution when interpreting these results.

According to the climate variability hypothesis organisms from more temperate environments are believed to have broader thermal tolerances compared to organisms from the tropics (Gutiérrez-Pesquera et al. 2016). This prediction originally stemmed from the observation that annual climatic variation is generally lower in the tropics compared to higher latitudes of temperate ecosystems. In that sense, temperate butterflies have developed various strategies to mitigate cooler and warmer conditions in temperate region such as migration to avoid extreme low temperatures such as monarch butterfly (Barve et al. 2012), darker coloration compared to those from warmer regions (Stelbrink et al. 2019), increased melanization in spring compared to summer periods (Kingsolver 1995), seasonal polyphenism to improve cold hardiness through overwintering (diapausing) (Van Dyck and Wiklund 2002) and thermal tolerance through summer aestivation such as *Maniola jurtina* (Grill et al. 2006).

Despite their complex ecological responses to climatic variability, body size is also changing over the years and shifts include both negative and, to a lesser extent, positive trends in Temperate Broadleaf and Mixed Forests. Global rising temperatures can justify the long decreasing trend we found between 1950 and 1990. Other anthropogenic stressors after World War II, such as urban expansion especially in Europe and North America and deforestation (Seto et al. 2011) transformed Earth’s land surface at unparalleled rates and scales directly affecting the distributions of many organisms (Newbold et al.

2016). The positive trends we observed, could be attributed to community reshuffling associated with urban-heat-island effect: this theory suggests that larger species, able to disperse and mitigate the low connectivity of ecological resources in urban settings (Cheptou et al. 2017), will slowly replace smaller body size species in urban settings (larger body size species fly in, smaller body size species fly out).

Acknowledging the factors associated with variability in insect morphology (Hawkins and Lawton 1995, Heidrich et al. 2021) brings novel insights into the species’ body size vulnerability to climate change. As an extension of this study, we plan to explore how much of the butterfly shrinkage we found can be attributed to direct effects of temperature and how much is indirectly filtering by other ecological traits (e.g., voltinism, first flight month, species temperature index) or other types of environmental filtering such as food availability and/or quality (Van Buskirk et al. 2010, Queiros et al. 2024).

## Code availability

The R-script used in the analysis is available at Zenodo (DOI: 10.5281/zenodo.14809535).

## Acknowledgements

We are grateful to Sevasti Lepenou for helping us with digitization in Goulandris Natural History Museum (GNHM) and Christina Kassara for helping us with spatial analysis.

## Author contributions

K.Z conceived the study. C.M.DM., M.C.M., M.R.L.W., and M.G. organized data use from ETHZ collection, K.Z., K.M.A., M.B., digitized butterfly collection of GNHM. K.Z., K.M.A., M.B., M.D., made identifications for GNHM collection. M.D. facilitated the data use from GNHM collection. K.Z., K.M.A., M.B., E.Kal., manually measured the specimens. Computer vision algorithm developed by T.S., P.S., with the help of K.Z. in customization. Data analysis conducted by K.Z., and G.C.A. Manuscript was originally drafted by K.Z., E.K., G.C.A., B.J.S., O.S., and V.K., with input from all the other authors.

