## Supporting Information for "Butterflies Are Shrinking: Evidence from the Past Century"

This file includes supplementary tables, figures, and additional analyses that support the conclusions of the main manuscript. It is intended to provide transparency and reproducibility for the reported findings.

**Contents:**

**Table S1-S7**

**Figure S1-S7**

**Body Size Sexual Dimorphism Section**: supplementary results and diagnostic plots for the first and second model both models.

**Aggregating Biomes Section**: supplementary results, plots and diagnostic plots for the second model.

For any questions regarding this material, please contact []

**Table S1.** We applied the global terrestrial biome map developed by Olson et al. (2001) in our dataset. Our dataset consisted of specimens from 13 biomes; three biomes had less than 20 specimens (*) and were excluded for further analyses.

| Biome number | Biome Description | Number of Individuals | Number of Species | From (Year) | Until (Year) | Years (total) | Years (total for1900-2010) |
| --- | --- | --- | --- | --- | --- | --- | --- |
| 1 | Tropical and Subtropical Moist Broadleaf Forests | 39 | 22 | 1908 | 2019 | 24 | 23 |
| 2 | Tropical and Subtropical Dry Broadleaf Forests | NA | NA | NA | NA | NA | NA |
| 3 | Tropical and Subtropical Coniferous Forests * | 2 | 1 | 1954 | 1967 | 2 | 2 |
| 4 | Temperate Broadleaf and Mixed Forests | 9197 | 303 | 1028 | 2017 | 151 | 104 |
| 5 | Temperate Coniferous Forests | 16500 | 251 | 1642 | 2017 | 150 | 105 |
| 6 | Boreal Forests/Taiga | 44 | 21 | 1878 | 1983 | 27 | 25 |
| 7 | Tropical and subtropical grasslands, savannas, and shrublands * | 1 | 1 | 1974 |  | 1 | 1 |
| 8 | Temperate Grasslands, Savannas, and Shrublands | 81 | 44 | 1893 | 2002 | 50 | 49 |
| 9 | Flooded Grasslands and Savannas | 22 | 10 | 1915 | 2002 | 8 | 8 |
| 10 | Montane Grasslands and Shrublands | 90 | 46 | 1900 | 2002 | 43 | 43 |
| 11 | Tundra | 24 | 12 | 1908 | 1982 | 15 | 15 |
| 12 | Mediterranean Forests, Woodlands, and Scrub | 5780 | 291 | 1872 | 2017 | 126 | 103 |
| 13 | Deserts and Xeric Shrublands | 137 | 54 | 1913 | 2002 | 60 | 60 |
| 98 | Lakes | NA | NA | NA | NA | NA | NA |
| 99 | Rock and Ice * | 3 | 1 | 1931 | 1982 | 3 | 3 |

**Table S2.** Output of our first model where forewing length is a function of three smoothers: a TPRS (thin plate regression spline) single common global smoother of year, a random effect of species and a nested random effect of species within years.

| Response variable | Parametric coefficients | Std. | Std.Error | t value | Pr(>\|t\|) |
| --- | --- | --- | --- | --- | --- |
| Wing length | (Intercept) | 2.88 | 0.11 | 25.61 | <0.001 |
|  | BIOME4 | -0.48 | 0.11 | -4.40 | <0.001 |
|  | BIOME5 | -0.47 | 0.11 | -4.33 | <0.001 |
|  | BIOME6 | -0.43 | 0.12 | -3.74 | <0.001 |
|  | BIOME8 | -0.50 | 0.11 | -4.41 | <0.001 |
|  | BIOME9 | -0.39 | 0.12 | -3.22 | <0.001 |
|  | BIOME10 | -0.41 | 0.11 | -3.68 | <0.001 |
|  | BIOME11 | -0.75 | 0.12 | -6.20 | <0.001 |
|  | BIOME12 | -0.43 | 0.11 | -3.95 | <0.001 |
|  | BIOME13 | -0.54 | 0.11 | -4.85 | <0.001 |
|  | Smooth terms | edf | Ref.df | F | p-value |
|  | s(Year.x) | 3.56 | 3.89 | 22.02 | <0.001 |
|  | s(GenusSpecies) | 261.26 | 578.00 | 998200.00 | <0.001 |
|  | s(GenusSpecies):Year.x | 321.54 | 578.00 | 1114000.00 | <0.001 |
| Explained deviance = 90.3% | |  |  |  |  |

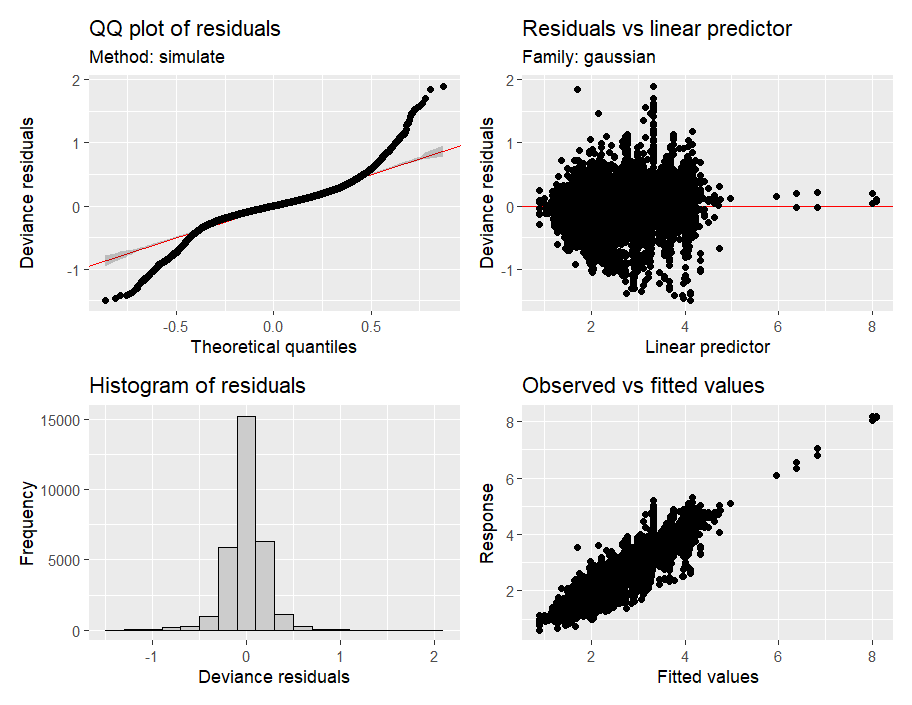

**Figure S1. Diagnostic plots for the first model**

**Table S3.** Output of the second model with a separate smoother per biome, a random effect of species and a nested random effect of species within years.

| Response variable | Parametric coefficients | Estimate | Std.Error | t value | Pr(>\|t\|) |
| --- | --- | --- | --- | --- | --- |
| Wing length | (Intercept) | 2.97 | 0.12 | 24.14 | <0.001 |
|  | BIOME4 | -0.59 | 0.12 | -4.84 | <0.001 |
|  | BIOME5 | -0.58 | 0.12 | -4.75 | <0.001 |
|  | BIOME6 | -0.58 | 0.13 | -4.48 | <0.001 |
|  | BIOME8 | -0.61 | 0.13 | -4.86 | <0.001 |
|  | BIOME9 | -0.46 | 0.26 | -1.73 | 0.083 |
|  | BIOME10 | -0.55 | 0.12 | -4.47 | <0.001 |
|  | BIOME11 | -1.01 | 0.15 | -6.81 | <0.001 |
|  | BIOME12 | -0.55 | 0.12 | -4.52 | <0.001 |
|  | BIOME13 | -0.66 | 0.12 | -5.34 | <0.001 |
|  | Smooth terms | edf | Ref.df | F | p-value |
|  | s(Year.x):BIOME1 | 3.40 | 3.66 | 8.89 | <0.001 |
|  | s(Year.x):BIOME4 | 3.90 | 3.99 | 52.57 | <0.001 |
|  | s(Year.x):BIOME5 | 1.80 | 2.24 | 1.38 | 0.261 |
|  | s(Year.x):BIOME6 | 2.04 | 2.49 | 1.89 | 0.136 |
|  | s(Year.x):BIOME8 | 1.58 | 1.94 | 0.92 | 0.330 |
|  | s(Year.x):BIOME9 | 1 | 1 | 0.04 | 0.849 |
|  | s(Year.x):BIOME10 | 1 | 1 | 0.10 | 0.753 |
|  | s(Year.x):BIOME11 | 2.04 | 2.38 | 3.31 | 0.032 |
|  | s(Year.x):BIOME12 | 3.72 | 3.95 | 10.36 | <0.001 |
|  | s(Year.x):BIOME13 | 1 | 1 | 10.29 | 0.001 |
|  | s(GenusSpecies) | 250.14 | 578 | 844400 | <0.001 |
|  | s(GenusSpecies):Year.x | 330.34 | 578 | 844900 | <0.001 |
| Explained deviance = 90.3% | |  |  |  |  |

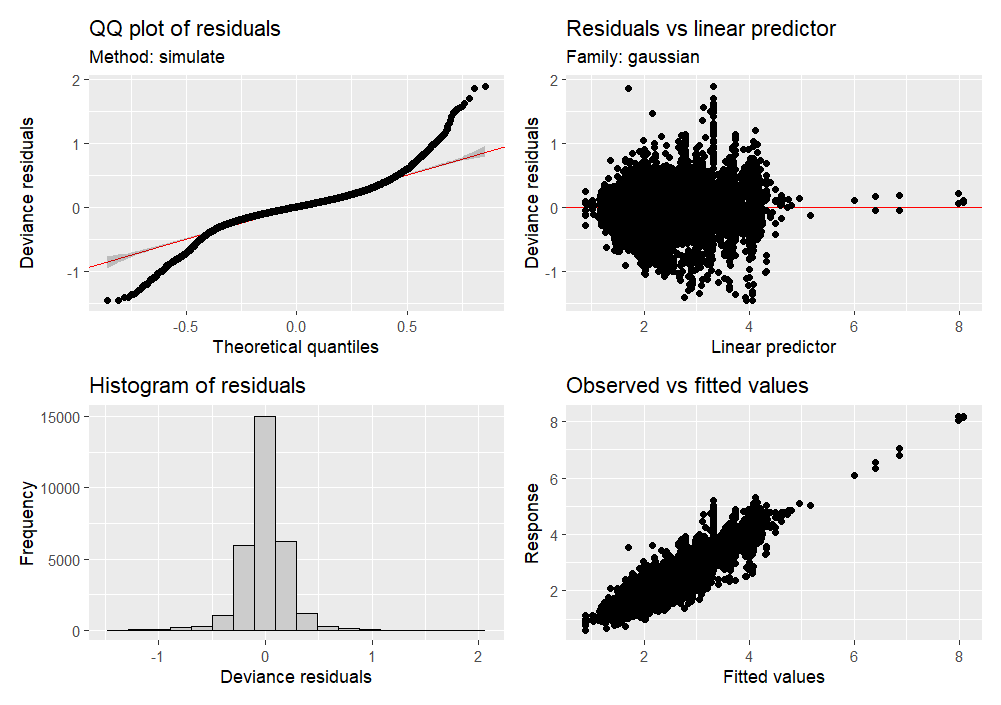

**Figure S2. Diagnostic plots for the second model.**

**Table S4.** Post-hoc comparisons of the contrasts of the parametric terms.

| Comparisons | Wald test | p-value |
| --- | --- | --- |
| BIOME 1 with 11 | 46.38 | <0.001 |
| BIOME 1 with 13 | 28.47 | <0.001 |
| BIOME 1 with 4 | 23.40 | <0.001 |
| BIOME 1 with 12 | 20.40 | <0.001 |
| BIOME 4 with 11 | 24.34 | <0.001 |
| BIOME 12 with 13 | 19.17 | <0.001 |
| BIOME 4 with 13 | 8.19 | 0.004 |
| BIOME 4 with 12 | 65.20 | <0.001 |
| BIOME 11 with 13 | 15.33 | <0.001 |
| BIOME 11 with 12 | 28.92 | <0.001 |

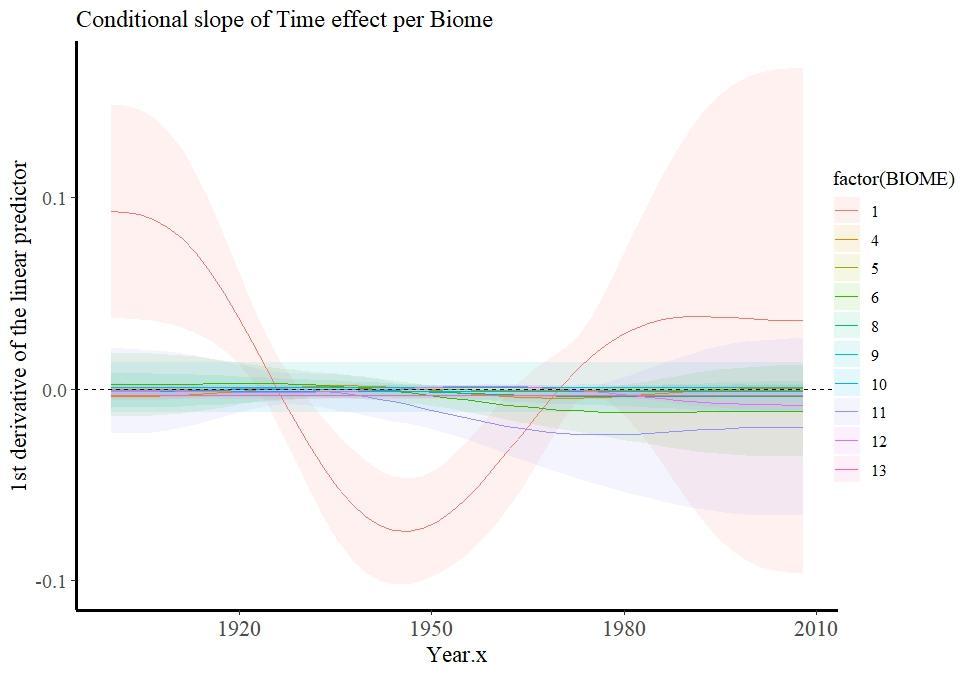

**Figure S3. Slopes of the fitted function over time as calculated by the first derivatives of smoother year per biome using *plot_slopes ()* function. It contains all 10 biomes included in the model. However, only biomes 1, 4, 11, 12, 13 were found to have a significant trend. Except for biome 1 and its pronounced negative and positive shifts (above and below zero), slopes of the studied biomes mainly remained below zero justifying the general decreasing trend for right forewing length we showed for the studied period.**

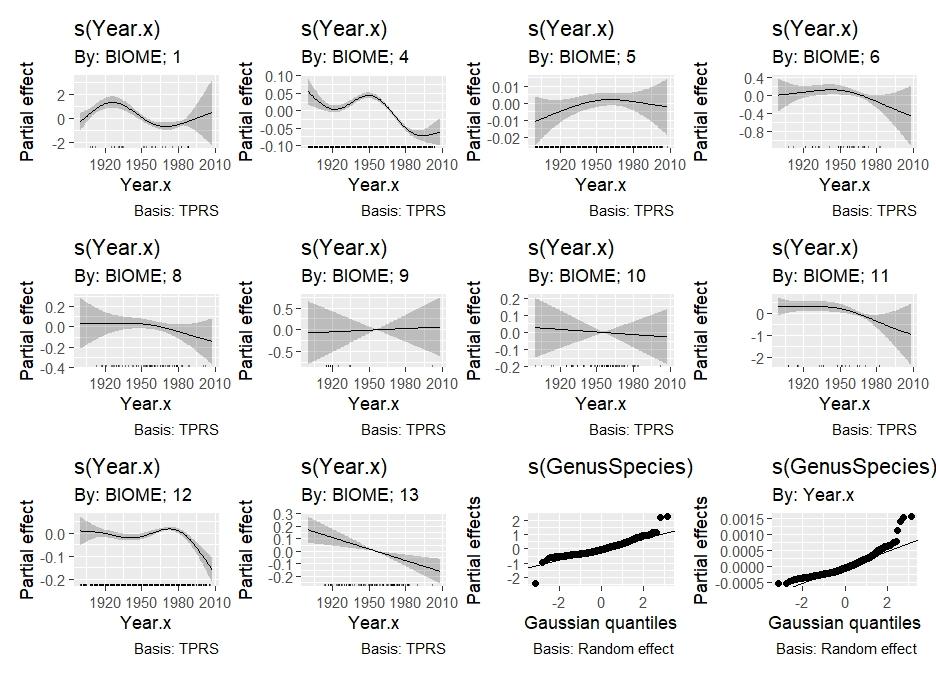

**Figure S4. Partial effects for all ten biomes we used in the second model. Note that trends were found to be significant for five biomes (1, 4, 11, 12, 13).**

**Body size sexual dimorphism section**

To account for sexual dimorphism effect, we rerun each model using only species for which the wing length is known to be the same between females and males. We used the European and Maghreb butterfly trait database (Middleton-Welling et al. 2020) and the morphological trait “Forewing length (FoL)” that corresponds to male and female average length with data obtained by various sources.

**Table S5.** Output of the first model considering only species for which wing length is considered to be equal between males and females. Forewing length is a function of three smoothers: a TPRS (thin plate regression spline) single common global smoother of year, a random effect of species and a nested random effect of species within years.

| Response variable | Parametric coefficients | Estimate | Std.Error | t value | Pr(>\|t\|) |
| --- | --- | --- | --- | --- | --- |
| Wing length | (Intercept) | 2.19 | 0.05 | 44.99 | <0.001 |
|  | BIOME5 | 0.01 | 0.00 | 2.76 | 0.005 |
|  | BIOME6 | 0.06 | 0.04 | 1.31 | 0.189 |
|  | BIOME8 | -0.07 | 0.04 | -1.93 | 0.053 |
|  | BIOME9 | 0.04 | 0.18 | 0.24 | 0.812 |
|  | BIOME10 | 0.00 | 0.11 | 0.02 | 0.985 |
|  | BIOME11 | -0.48 | 0.09 | -5.58 | <0.001 |
|  | BIOME12 | -0.01 | 0.01 | -2.05 | 0.04 |
|  | BIOME13 | -0.31 | 0.04 | -7.34 | <0.001 |
|  | Smooth terms | edf | Ref.df | F | p-value |
|  | s(Year.x) | 3.70 | 3.95 | 18.77 | <0.001 |
|  | s(GenusSpecies) | 83.47 | 202.00 | 134811.95 | 0.708 |
|  | s(GenusSpecies):Year.x | 121.17 | 202.00 | 341862.08 | 0.004 |
| Explained deviance = 90.9% | |  |  |  |  |

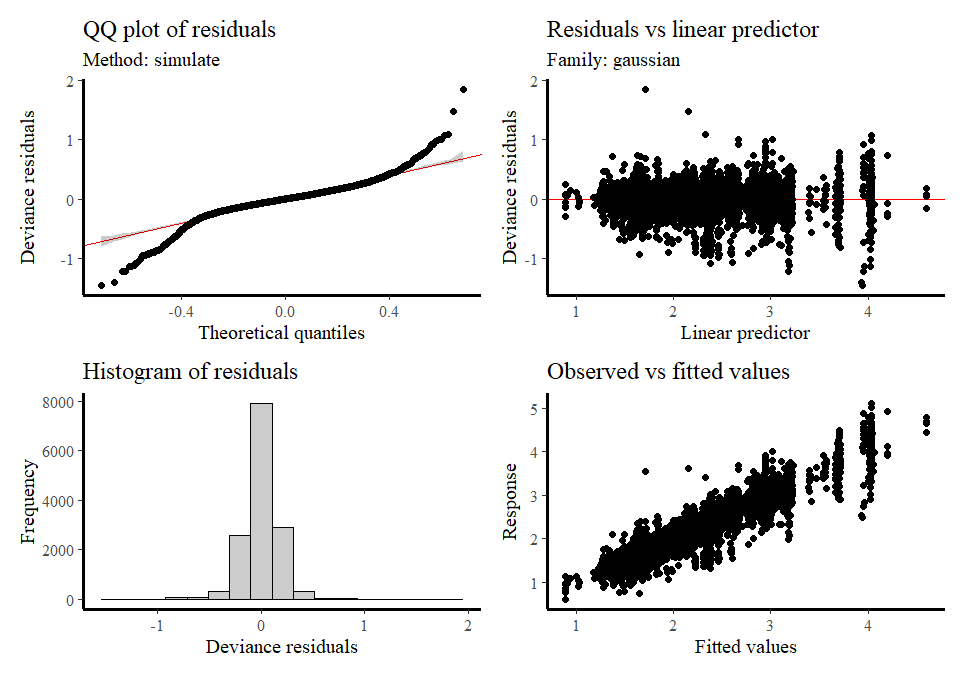

**Figure S5. Diagnostic plots for the second model.**

**Table S6.** Output of the second model considering only species for which wing length is considered to be equal between males and females. Model considers a separate smoother per biome, a random effect of species and a nested random effect of species within years.

| Response variable | Parametric coefficients | Estimate | Std.Error | t value | Pr(>\|t\|) |
| --- | --- | --- | --- | --- | --- |
| Wing length | (Intercept) | 2.18 | 0.05 | 44.48 | <0.001 |
|  | BIOME5 | 0.02 | 0.00 | 3.44 | <0.001 |
|  | BIOME6 | -0.01 | 0.07 | -0.21 | 0.836 |
|  | BIOME8 | -0.07 | 0.04 | -1.83 | 0.067 |
|  | BIOME9 | 0.00 | 0.00 | NaN | NaN |
|  | BIOME10 | -0.32 | 0.32 | -1.02 | 0.309 |
|  | BIOME11 | -0.39 | 0.13 | -3.07 | 0.002 |
|  | BIOME12 | -0.01 | 0.01 | -2.31 | 0.021 |
|  | BIOME13 | -0.34 | 0.04 | -7.88 | <0.001 |
|  | Smooth terms | edf | Ref.df | F | p-value |
|  | s(Year.x):BIOME4 | 3.78 | 3.97 | 35.47 | <0.001 |
|  | s(Year.x):BIOME5 | 1.00 | 1.00 | 0.01 | 0.912 |
|  | s(Year.x):BIOME6 | 1.50 | 1.80 | 1.49 | 0.218 |
|  | s(Year.x):BIOME8 | 1.91 | 2.34 | 1.62 | 0.227 |
|  | s(Year.x):BIOME9 | 1.00 | 1.00 | 0.07 | 0.798 |
|  | s(Year.x):BIOME10 | 1.00 | 1.00 | 1.13 | 0.288 |
|  | s(Year.x):BIOME11 | 1.66 | 1.79 | 2.55 | 0.046 |
|  | s(Year.x):BIOME12 | 3.30 | 3.73 | 10.21 | <0.001 |
|  | s(Year.x):BIOME13 | 1.00 | 1.00 | 14.44 | <0.001 |
|  | s(GenusSpecies) | 71.02 | 202 | 87930 | 0.854 |
|  | s(GenusSpecies):Year.x | 132.46 | 202 | 239500 | 0.005 |
| Explained deviance = 91% | |  |  |  |  |

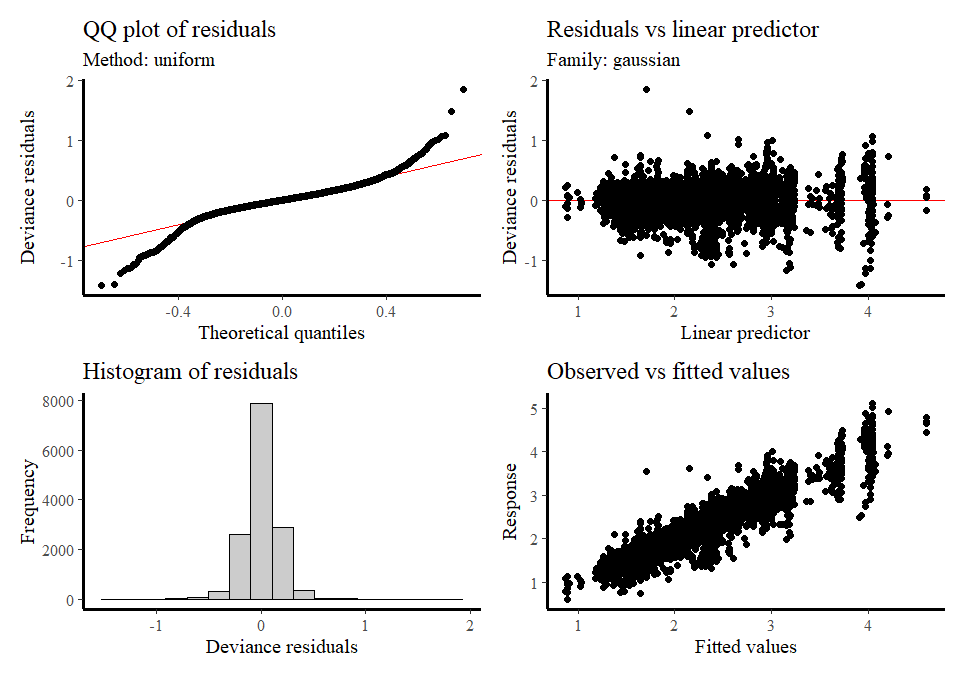

**Figure S6. Diagnostic plots for the second model.**

**Aggregating biomes section**

We aggregated the temperate (biomes 4; 5; 8) and tropical (biomes 1;3;7) biomes and we rerun the second model. Below is the output, plots of biomes and diagnostic plots.

**Table S7.** Output of the second model where temperate biomes (4, 5, 8) and tropical biomes (1, 3, 7) were combined. Model uses a separate smoother per biome, a random effect of species and a nested random effect of species within years.

| Response variable | Parametric coefficients | Estimate | Std.Error | t value | Pr(>\|t\|) |
| --- | --- | --- | --- | --- | --- |
| Wing length | (Intercept) | 2.40 | 0.06 | 41.27 | <0.001 |
|  | BIOME 9 | 0.11 | 0.24 | 0.46 | 0.648 |
|  | BIOME 10 | 0.03 | 0.07 | 0.42 | 0.672 |
|  | BIOME 11 | -0.43 | 0.10 | -4.49 | <0.001 |
|  | BIOME 12 | 0.03 | 0.05 | 0.63 | 0.527 |
|  | BIOME 13 | -0.08 | 0.05 | -1.62 | 0.106 |
|  | BIOME 1_3_7 | 0.56 | 0.11 | 5.01 | <0.001 |
|  | BIOME 4_5_8 | -0.01 | 0.04 | -0.17 | 0.866 |
|  | Smooth terms | edf | Ref.df | F | p-value |
|  | s(Year.x):BIOME 6 | 2.06 | 2.51 | 1.84 | 0.144 |
|  | s(Year.x):BIOME 9 | 1.00 | 1.00 | 0.01 | 0.918 |
|  | s(Year.x):BIOME 10 | 1.00 | 1.00 | 0.06 | 0.812 |
|  | s(Year.x):BIOME 11 | 2.01 | 2.35 | 3.37 | 0.031 |
|  | s(Year.x):BIOME 12 | 3.71 | 3.95 | 9.64 | <0.001 |
|  | s(Year.x):BIOME 13 | 1.00 | 1.00 | 10.19 | 0.001 |
|  | s(Year.x):BIOME 1_3_7 | 3.58 | 3.81 | 9.92 | <0.001 |
|  | s(Year.x):BIOME 4_5_8 | 3.76 | 3.97 | 26.34 | <0.001 |
|  | s(GenusSpecies) | 566.43 | 579.00 | 442.61 | <0.001 |
| Explained deviance = 90.2% | |  |  |  |  |

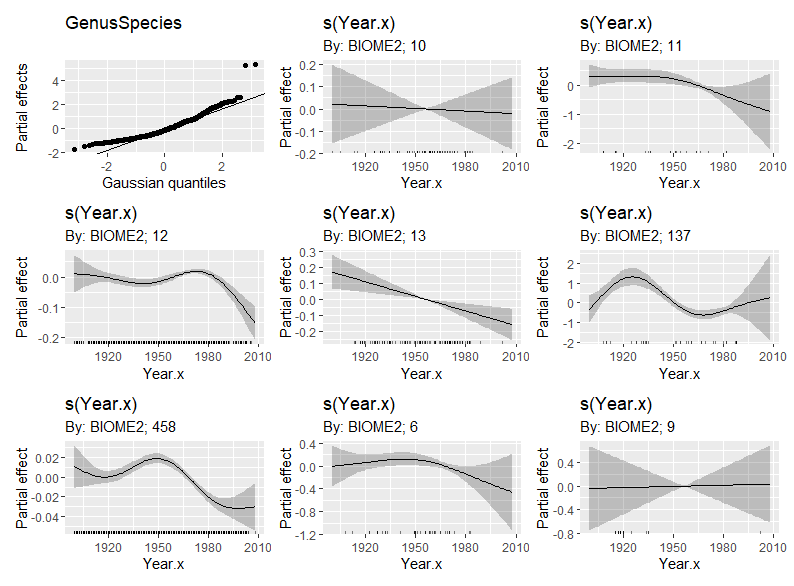

**Figure S7. Partial effects for eight biomes we used in the second model after aggregating the temperate and tropical biomes.**
